# Accurate quantification of copy-number aberrations and whole-genome duplications in multi-sample tumor sequencing data

**DOI:** 10.1101/496174

**Authors:** Simone Zaccaria, Benjamin J. Raphael

## Abstract

Copy-number aberrations (CNAs) and whole-genome duplications (WGDs) are frequent somatic mutations in cancer. Accurate quantification of these mutations from DNA sequencing of bulk tumor samples is complicated by varying tumor purity, admixture of multiple tumor clones with distinct mutations, and high aneuploidy. Standard methods for CNA inference analyze tumor samples individually, but recently DNA sequencing of multiple samples from a cancer patient – e.g. from multiple regions of a primary tumor, matched primary/metastases, or multiple time points – has become common. We introduce a new algorithm, <u>H</u>olistic <u>A</u>llele-specific <u>T</u>umor <u>Co</u>py-number <u>He</u>terogeneity (HATCHet), that infers allele and clone-specific CNAs and WGDs *jointly* across multiple tumor samples from the same patient, and that leverages the relationships between clones in these samples. HATCHet provides a fresh perspective on CNA inference and includes several algorithmic innovations that overcome the limitations of existing methods, resulting in a more robust approach even for single-sample analysis. We also develop MASCoTE (<u>M</u>ultiple <u>A</u>llele-specific <u>S</u>imulation of <u>Co</u>py-number <u>T</u>umor <u>E</u>volution), a framework for generating realistic simulated multi-sample DNA sequencing data with appropriate corrections for the differences in genome lengths between the normal and tumor clone(s) present in mixed samples. HATCHet outperforms current state-of-the-art methods on 256 simulated tumor samples from 64 patients, half with WGD. HATCHet’s analysis of 49 primary tumor and metastasis samples from 10 prostate cancer patients reveals subclonal CNAs in only 29 of these samples, compared to the published reports of extensive subclonal CNAs in all samples. HATCHet’s inferred CNAs are also more consistent with the reports of polyclonal origin and limited heterogeneity of metastasis in a subset of patients. HATCHet’s analysis of 35 primary tumor and metastasis samples from 4 pancreas cancer patients reveals subclonal CNAs in 20 samples, WGDs in 3 patients, and tumor subclones that are shared across primary and metastases samples from the same patient – none of which were described in published analysis of this data. HATCHet substantially improves the analysis of CNAs and WGDs, leading to more reliable studies of tumor evolution in primary tumors and metastases.

## 1 Introduction

Cancer results from the accumulation of somatic mutations in cells, yielding a heterogeneous tumor composed of distinct subpopulations of cells, or *clones*, with different complements of mutations^1^. Quantifying this intra-tumor heterogeneity and inferring tumor evolution have been shown to be crucial in cancer treatment and prognosis^2–4^. Copy-Number Aberrations (CNAs) are frequent somatic mutations in cancer that amplify or delete one or both the alleles of genomic segments, chromosome arms, or even entire chromosomes^5^. In addition whole-genome duplication (WGD), a doubling of all chromosomes, is a frequent event in cancer with an estimated frequency higher than 30% in recent pan-cancer studies^5–8^. Accurate inference of CNAs and WGDs is crucial for quantifying intra-tumor heterogeneity and reconstructing tumor evolution, even when analyzing only single-nucleotide variants (SNVs)^9–13^.

In principle, CNAs can be detected in DNA sequencing data by examining differences between the observed and expected counts of sequencing reads that align to a locus, quantified either by the read-depth ratio (RDR) of genomic segments or by the B-allele frequency (BAF) of heterozygous germline SNPs. In practice, the inference of CNAs and WGDs from DNA sequencing data is challenging, particularly for bulk tumor samples that are mixtures of thousands-millions of cells. In such a mixture the signal from the observed reads is a superposition of the signals from normal and cancer cells, with the cancer cells further divided into one or more *clones*. One thus needs to *deconvolve*, or separate, this mixed signal into the individual components arising from each of these clones. This deconvolution is complicated as both the CNAs and the proportion of cells originating from each clone in the mixture are unknown; in general the deconvolution problem is *underdetermined* with multiple equivalent solutions. In the past few years, over a dozen methods have been developed to solve different simplified versions of this copy-number deconvolution problem^6,9,14–27^. These methods rely on various simplifying assumptions such as only one tumor clone is present in the mixture, WGDs are not present, etc. While these assumptions remove ambiguity in copy-number deconvolution, it is not clear that the resulting solutions are accurate, particularly in cases of highly aneuploid tumors.

While single-cell DNA sequencing^28^ obviates the need for copy-number deconvolution, it remains a specialized technique with various technical and financial challenges, and thus is not yet widely-used in sequencing of cancer patients, particularly in clinical settings. A valuable intermediate between DNA sequencing of single cells and DNA sequencing of a single bulk tumor sample is DNA sequencing of multiple tumor samples from the same patient – including multiple regions of a primary tumor, matched primary and metastases, or longitudinal samples^11,12,26,29–31^. A number of approaches have demonstrated that *simultaneous* analysis of SNVs from multiple samples from the same patient helps resolve uncertainties in clustering SNVs into clones^32,33^ and reduces ambiguities in inferring phylogenetic trees^11,29,34–36^. Remarkably, with one exception^25^, available methods for inferring CNAs analyze *individual* samples, losing the important information that multiple samples from the same patient are correlated via the shared evolutionary process that gave rise to the tumor.

To slice through the thicket of uncertainty in copy-number deconvolution, we introduce <u>H</u>olistic <u>A</u>llele-specific <u>T</u>umor <u>Co</u>py-number <u>He</u>terogeneity (HATCHet), an algorithm that infers allele and clone-specific CNAs as well as the proportions of distinct tumor clones *jointly* across one or more samples from the same patient. HATCHet provides a fresh perspective on CNA inference and includes several algorithmic innovations that overcome the limitations of existing methods. First, HATCHet solves a simultaneous matrix factorization problem which models allele-specific copy numbers, the dependencies *between genomic segments across clones*, and the dependencies *between clones across samples*. In contrast, existing methods do not infer allele-specific copy numbers^20–24^, consider each segment independently^6,9,14–19^, do not preserve clonal structure across samples^21,22,26,27^, or assume all samples comprise the same set of few clones^25^. Second, HATCHet *globally* clusters RDR and BAF jointly along the genome and across all samples, while existing methods rely on *local* clustering of neighboring loci. Third, HATCHet performs the copy-number deconvolution in the natural coordinates of integer copy numbers and clone proportions. In contrast, existing methods use parameters of *tumor purity* and *tumor ploidy* (or equivalent parameters) to select among different solutions of the copy-number deconvolution problem. However, tumor purity and ploidy are *composite* parameters that average over the unknown copy numbers and proportions, and such averages may not adequately distinguish between multiple equally-plausible solutions. Last, HATCHet defines a model selection criterion that evaluates the trade-off between the inference of multiple tumor clones in a sample versus the inference of a WGD. In contrast, existing methods exclude WGD from model selection, do not consider this trade-off, and/or evaluate solutions in the composite parameters of tumor purity and ploidy that are ill-suited to model selection.

We compare HATCHet with 4 current state-of-the-art methods: Battenberg^9^, TITAN^17^, THetA^21,22^, and cloneHD^25^ on a simulated dataset comprising 256 samples from 64 patients, half with WGD. Since current approaches to simulate tumor sequencing data incorrectly assume that all genomes in the mixture have the same length^15–17,25,37–42^, we also develop MASCoTE (<u>M</u>ultiple <u>A</u>llele-specific <u>S</u>imulation of <u>Co</u>py-number <u>T</u>umor <u>E</u>volution), a simulation framework to generate sequencing reads with appropriately corrections for the differences in genome lengths between the normal and tumor clone(s) present in multiple mixed samples. We show that HATCHet outperforms all the other methods (and the consensus of these methods) in the inference of CNAs, their proportions, and WGDs – even when using only single samples.

We apply HATCHet to analyze CNAs in two published whole-genome, multi-sample tumor sequencing datasets. On a dataset of 49 samples of primary tumors and metastases from 10 prostate cancer patients^11^, HATCHet produces copy numbers and proportions which significantly simplify the extensive subclonality reported in published analysis while better explaining the data. Published analysis reports subclonal CNAs – CNAs that are present in only a subset of the tumor cells in a sample – in all 49 samples, while HATCHet identifies 20 samples without subclonal CNAs and also yields more consistent predictions of WGDs. Moreover, HATCHet identifies CNA-derived clones that are shared among distinct metastases consistent with previous reports of polyclonal origin or limited heterogeneity. On a dataset of 35 primary tumor samples and metastases from 4 pancreas cancer patients^30^, HATCHet identifies subclonal CNAs in 20 samples and WGDs in 3 patients while published analysis excluded the presence of subclonal CNAs and WGDs. HATCHet also identifies tumor subclones that are shared across primary and metastases samples from the same patient. Finally, we show that the observed read counts of somatic SNVs are more consistent with the CNAs inferred by HATCHet than CNAs inferred by other methods. Thus, HATCHet substantially improves the analysis of CNAs and WGDs, leading to more reliable studies of tumor evolution in primary tumors and metastases.

## 2 Results

### 2.1 <u>H</u>olistic <u>A</u>llele-specific <u>T</u>umor <u>Co</u>py-number <u>He</u>terogeneity (HATCHet) algorithm

Suppose that we sequence DNA from *k* bulk-tumor samples from the same patient. We assume that each sample is a mixture of at most *n* clones, including the normal diploid clone and one or more tumor clones which are distinguished by CNAs (Fig. 1A). Each CNA alters the number of copies of a contiguous genomic region from one of the two homologous chromosomes, which define the two alleles of each region. Thus, we represent the accumulation of CNAs in all clones by partitioning the *L* genomic positions of the reference genome into *m* segments, with each segment *s* consisting of *ℓ_s_* neighboring positions with the same copy number in a clone. We model the pair of allele-specific copy numbers of segment *s* in a clone *i* as a *copy-number state* (*a*_*s*,*i*_, *b*_*s*, *i*_). We represent the allele and clone-specific copy numbers of all clones by two *m* × *n* matrices *A* = [*a*_*s*,*i*_] and *B* = [*b*_*s*,*i*_].

**Fig. 1:**
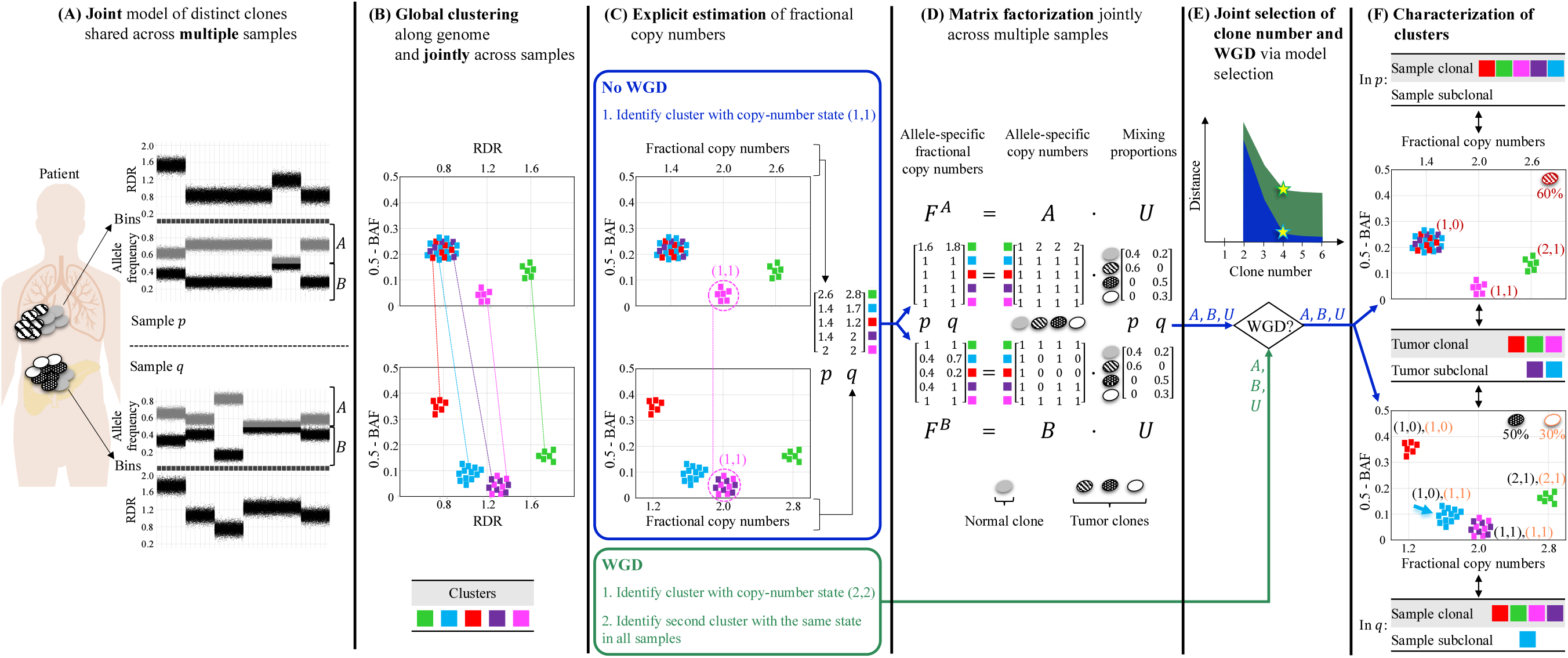
Overview of <u>H</u>olistic <u>A</u>llele-specific <u>T</u>umor <u>C</u>opy-number <u>He</u>terogeneity (HATCHet) algorithm. (A) HATCHet analyzes the read-depth ratio (RDR) and the B-allele frequency (BAF) in bins of the reference genome (black squares) jointly from multiple tumor samples. Here, we show two tumor samples *p* and *q*. (B) HATCHet globally clusters the bins based on RDR and BAF along the entire genome and jointly across samples *p* and *q*. Each cluster (color) includes bins with the same copy-number state within each clone present in *p* or *q*. (C) HATCHet estimates the fractional copy number of each cluster. If there is no WGD, the identification of the cluster (magenta) with copy-number state (1, 1) is sufficient and RDRs are scaled correspondingly. If a WGD occurs, HATCHet finds the cluster with copy-number state (2,2) (same magenta cluster) and a second cluster having an identical copy-number state in all tumor clones. (D) HATCHet factorizes the allele-specific fractional copy numbers *F^A^*, *F^B^* into the allele-specific copy numbers **A*, *B**, respectively, and the clone proportions *U*. Here there is a normal clone and 3 tumor clones. (E) HATCHe’s model selection criterion identifies the matrices **A*, *B** and *U* in the factorization while evaluating the fit according to both the inferred number of clones and presence/absence of a WGD. (F) Clusters are classified by their inferred copy-number states in each sample. *Sample-clonal clusters* have a unique copy-number state in the sample and correspond to evenly-spaced positions in the scaled RDR-BAF plot (vertical grid lines in each plot). *Sample-subclonal clusters* (e.g. cyan in *p*) have different copy-number states in a sample and thus correspond to intermediate positions in the scaled RDR-BAF plot. *Tumor-clonal clusters* have identical copy-number states in all tumor clones – thus they are sample-clonal clusters in every sample and preserve their relative positions in scaled-RDR-BAF plots. In contrast, tumor-subclonal clusters have different copy-number states in different tumor clones and their relative positions in the scaled RDR-BAF plot varies across samples (e.g. purple cluster).

DNA sequencing data from the *k* samples does not directly measure *A* and *B*, but rather measures a *mixture* of copy number states. In particular, for each segment *s* and each sample *p*, we observe two allele-specific *fractional copy numbers* 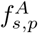 and 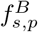. The fractional copy numbers across the *m* segments for *k* samples form two *m* × *k* matrices *F^A^* and *F^B^*. Tumor samples from the same patient are related by the somatic evolutionary process that gave rise to the tumor. Specifically, let the clone proportion *u*_*i*,*p*_ be the fraction of cells in sample *p* that belong to clone *i*. We represent the clone proportions for all clones and samples by an *n* × *k* matrix *U* = [*u*_*i*,*p*_]. As such, the *fractional copy numbers* are determined by the allele-specific copy numbers and clone proportions via the equations *F^A^* = *AU* and *F^B^* = *BU*.

The copy-number deconvolution problem corresponds to the problem of *simultaneously* factoring *F^A^* and *F^B^* into the corresponding allele-specific copy numbers *A*, *B* and clone proportions *U*. This matrix-factorization formulation differs in three key ways from current approaches to CNA inference. First, we model allele-specific copy numbers, while many existing methods do not^20–24^. Second, we model dependencies *between samples* while all current approaches (except^25^) analyze samples independently. Third, we model dependencies *between segments* as clones, while all other widely-used methods either consider each segment independently^6,9,17–19^ (Fig. S1), do not preserve clonal structure across samples^21,22,26,27^, or assume all samples comprise the same set of few clones^25^.

While our simultaneous matrix factorization is a mathematically elegant description of the joint copy-number deconvolution of multiple samples, there are several practical issues that must be addressed to derive a useful algorithm for DNA sequencing data: (1) segments are unknown; (2) *F^A^* and *F^B^* are not directly observed; (3) measurement errors in *F^A^* and *F^B^* may affect the existence of factorizations *F^A^* = *AU* and *F^B^* = *BU*; (4) multiple factorizations leading to degenerate solutions may exist; (5) the number *n* of clones and the occurrence of WGD are unknown a priori. We develop the algorithm HATCHet (<u>H</u>olistic <u>A</u>llele-specific <u>T</u>umor <u>Co</u>py-number <u>He</u>terogeneity) to address these issues.

First, we design a global clustering approach to infer the segments that underwent CNAs. All existing methods for CNA inference rely on local clustering of neighboring genomic regions with similar values of RDR and/or BAF (Fig. S2). In contrast, HATCHet globally clusters genomic regions *along* the genome and *jointly across* multiple samples (Fig. 1B).

Second, we introduce a rigorous criterion to explicitly estimate the fractional copy numbers *F^A^* and *F^B^* from DNA sequencing data, Nearly all methods – including widely-used methods such as ABSOLUTE^6^, ASCAT^14^, Battenberg^9^, TITAN^17^, cloneHD^25^, and others^15,16,18–20,23,24,26,27^ do not attempt to directly infer fractional copy numbers, but rather attempt to fit the parameters of tumor ploidy and purity (or equivalent parameters). However, these are *composite* parameters that sum the contributions of the *unknown* copy numbers and proportions of multiple clones. The dependency between the values of these parameters and the underlying clonal composition of the mixed tumor sample is complicated to model and challenging to infer^21,22,25^. The consequence is that tumor ploidy and purity are not good coordinates to evaluate tumor mixtures as many different clonal compositions may be equally plausible in these coordinates, particularly when more than one tumor clone is present or a WGD occurs (Fig. S3, Fig. S4, and Fig. S5). Not surprisingly, manual inspection of the results from current methods is often required to evaluate the presence of WGD^6,7,12^. In contrast, HATCHet explicitly infers *F^A^* and *F^B^* using a mathematical result which states that the identification of the copy-number state for one cluster (or two clusters in the case of WGD) is sufficient to scale observed read counts into fractional copy numbers without any further information about the tumor composition (Fig. 1C).

Last, we deploy an explicit model-selection criterion to address the three remaining issues. We address measurement errors in *F^A^* and *F^B^* by minimizing the differences |*F^A^* − *AU* | and |*F^B^* − *BU* |, allowing for cases where an exact factorization does not exist. We address the issue of multiple equivalent solutions by including several reasonable constraints in the allowed factorizations including a maximum copy number (*a*_*s*,*i*_ + *b*_*s*,*i*_ ≤ *c*_max_), a minimum clone proportion (either *u*_*i*,*p*_ ≥ *u*_min_ or *u*_*i*,*p*_ = 0), and enforcing evolutionary relationship among the tumor clones. HATCHet solves the resulting optimization problem using a coordinate-descent algorithm (Fig. 1D). Finally, HATCHet jointly infers the number of clones *n* and predicts the presence/absence of a WGD using a model-selection criterion that explicitly models the trade-off between subclonal CNAs (resulting from higher total number of clones and more clones present in a sample) and WGD (Fig. 1E). In contrast, existing methods ignore this trade-off and do not include WGD (modeled as higher values of tumor ploidy) in the model selection^6,9,14–22,25–27^. Our model-selection criterion uses the natural coordinates of allele-specific copy numbers (matrices *A*, *B*) and clone proportions (matrix *U*), instead of the composite parameters tumor purity and tumor ploidy that average over the clonal composition of a sample.

Further details of HATCHet are in Section 4.

### 2.2 HATCHet outperforms existing methods for copy-number deconvolution

We compare HATCHet with four current state-of-the-art methods, Battenberg^9^, TITAN^17^, THetA^21,22^, and cloneHD^25^, on simulated data. The simulation of DNA sequencing data from bulk tumor samples containing large-scale CNAs is not straightforward, and subtle mistakes are common in previously published studies. Most current studies that simulate sequencing reads from mixed samples do not account for the different genome lengths of distinct clones^15–17,25,37–42^, and this leads to incorrect simulation of read counts (Fig. S6 and Fig. S7). Therefore, we develop a new simulation framework MASCoTE (<u>M</u>ultiple <u>A</u>llele-specific <u>S</u>imulation of <u>Co</u>py-number <u>T</u>umor <u>E</u>volution) to simulate the genomes of clones with distinct CNAs and WGDs, and to correctly generate multi-sample bulk tumor sequencing data (Fig. S8 and details in Section 4). We simulate DNA sequencing reads from 256 samples (2–3 tumor clones per sample) for 64 patients (3-5 samples per patient), half with a WGD and half without a WGD (Fig. S18). Further details regarding the simulations, the experimental setup for all methods, additional results, and a complete description of all the comparison metrics are in Supplementary Note C.1.

To assess the inference of CNAs and their proportions, we first run all methods on the 128 samples from 32 patients without a WGD and also provide the true value of the main parameters (e.g. tumor ploidy, number of clones, and maximum copy number) required for each method. This provides a baseline comparison of the performance of each method in determining copy numbers and proportions. We find that HATCHet outperforms all other methods, even when we run HATCHet on individual samples without taking advantage of HATCHet’s joint inference across multiple samples (Fig. 2A, Fig. S19–S22). The gain on single-samples is likely due to HATCHet’s other key features described above. In particular, we observe that Battenberg and TITAN, which infer the copy numbers of genomic regions independently, perform significantly worse than THetA, cloneHD, and HATCHet, which group copy numbers into the clones present in a sample. Furthermore, we observe that cloneHD – the only existing method that considers multiple samples simultaneously – shows only a modest gain over THetA which analyzes samples individually; moreover cloneHD performs worse than single-sample HATCHet. This suggests that cloneHD is not deriving maximum benefit from multiple samples, perhaps because its model assumes that the same few clones are present in all samples.

**Fig. 2:**
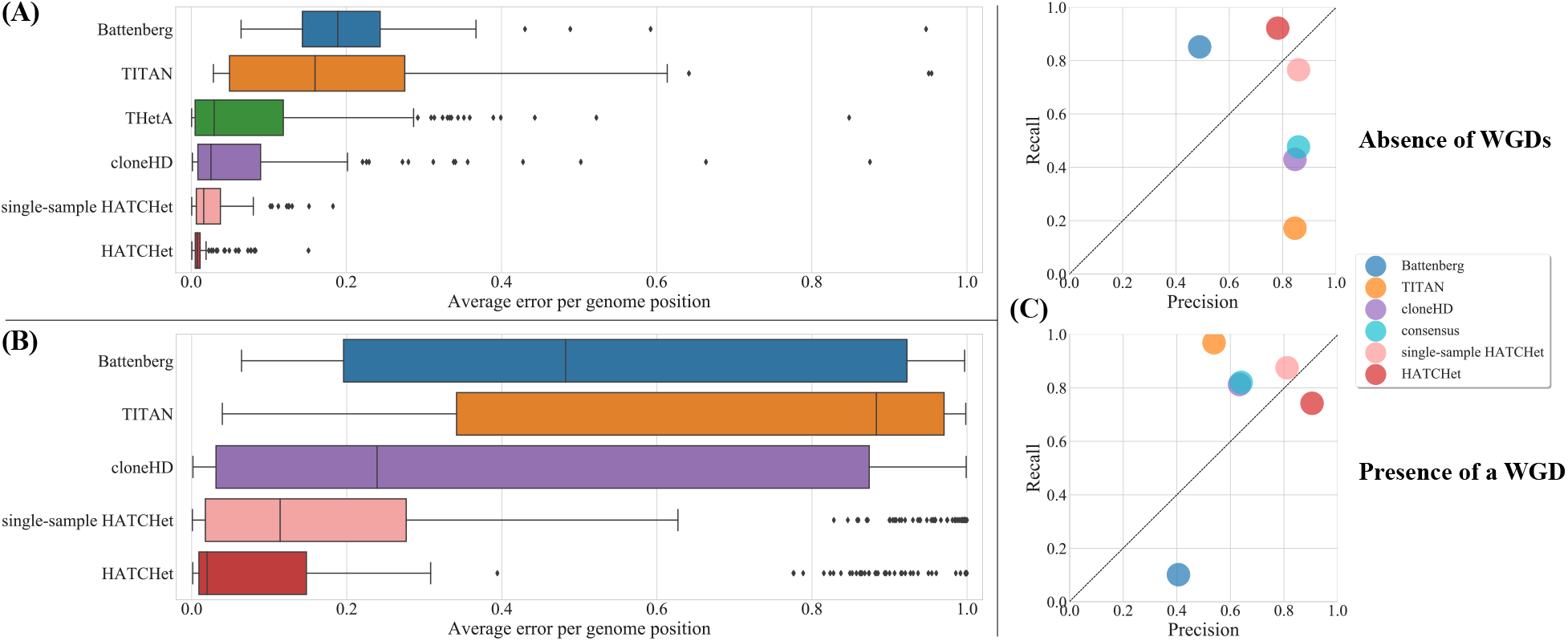
HATCHet outperforms existing methods in the inference of copy-number aberrations, their proportions, and whole-genome duplications. (A) Average error per genome position for the copy-number states and their proportions inferred by each method on 128 simulated tumor samples from 32 patients without a WGD, and where each method was provided with the true values of the main parameters (e.g. tumor ploidy, number of clones, and maximum copy number). HATCHet outperforms all the other methods even when it considers single samples individually (single-sample HATCHet). (B) Average error per genome position on 256 simulated samples from 64 patients, half with a WGD, and where each method infers all relevant parameters including tumor ploidy, number of clones, etc. HATCHet outperforms all the other methods, even when considering single samples individually (single-sample HATCHet). (C) Average precision and recall in the prediction of the absence of a WGD and the presence of a WGD in a sample. HATCHet is the only method with high precision and recall (> 75%) in both the cases, even compared to a consensus of the other methods based on a prediction for majority. While Battenberg underestimates the presence of WGDs (< 20% recall), TITAN and cloneHD overestimates the absence of WGDs (< 20% and < 50% recall, respectively).

To assess the simultaneous prediction of WGD and inference of CNAs and proportions, we next run the methods on all the 256 samples from all 64 patients, requiring that each method infers all relevant parameters including tumor ploidy, number of clones, etc. We set the maximum copy number of a segment to 8, and excluded THetA from this comparison as it does not automatically infer presence/absence of WGDs. Not surprisingly, in this more challenging setting the performance of all methods is lower, but HATCHet continues to outperform the other methods – even when considering single samples individually (Fig. 2B and Fig. S23–S27), and even when assessing the prediction of amplified/deleted segments independently from the presence of a WGD (Fig. S28). HATCHet is the only method with high (> 75%) precision and recall in the identification of both the presence and absence of a WGD, while other methods are biased towards one of the two predictions (Fig. 2C and Fig. S29). We observe the same bias even when taking the consensus of the other methods, as was done in the recent PCAWG analysis of > 2500 whole-cancer genomes^7^ (Fig. 2C). The significantly lower performance of all existing methods relative to the previous comparison as well as the high error-rate in the prediction of WGDs illustrate the challenges in selecting a solution using the coordinates of tumor purity and ploidy (further details in Section 4.4). While these coordinates are used by all these existing methods, HATCHet’s model-selection criterion is based on the natural variables of the problem, enabling HATCHet to achieve robust performance.

### 2.3 HATCHet identifies well-supported subclonal CNAs

We use HATCHet to analyze two published whole-genome, multi-sample tumor sequencing datasets: 49 primary and metastatic tumor samples from 10 metastatic prostate cancer patients^11^ and 39 primary and metastatic tumor samples from 4 pancreatic cancer patients^30^. Although both of these datasets contained multiple samples from the same patient, the published analyses inferred CNAs in each sample independently. Moreover, these studies reached opposite conclusions regarding the landscape of CNAs in these tumors. Gundem et al.^11^ report *subclonal CNAs* – CNAs present in only a subset of the tumor cells in a sample – in *all* primary and metastatic prostate samples. In contrast, Makohon-Moore et al.^30^ report *no* subclonal CNAs in the primary and metastatic pancreatic samples. An important question is whether this difference is a result of cancer-type specific or patient-specific differences in CNA evolution of these tumors, or possibly a result of analytical differences as Battenberg^9^ was used to infer CNAs in the prostate cancer publication^11^ and Control-FREEC^20^ was used to infer CNAs in the pancreas cancer publication^30^.

In the prostate cancer dataset, HATCHet identifies subclonal CNAs in 29 samples, while Battenberg identifies subclonal CNAs in all 49 samples (Fig. 3A). In the 29 samples HATCHet and Battenberg report similar fractions of the genome with subclonal CNAs (Fig. 3B and Fig. S30B). Within a sample, clonal CNAs correspond to *sample-clonal* clusters of genomic regions that have the same copy-number state in all tumor clones, while subclonal CNAs correspond to *sample-subclonal* clusters of genomic regions that have values of RDR and BAF that are intermediate between those of sample-clonal clusters (Fig. 1F and Fig. S9). We find that in each of the 29 samples where Battenberg and HATCHet report suclonal CNAs, there are clear sample-subclonal clusters with RDR and BAF values that are clearly distinct and intermediate between those of sample-clonal clusters (Fig. 3C). Moreover, these sample-subclonal clusters correspond to large genomic regions (Fig. 3D). In contrast, in each of the remaining 20 samples where only Battenberg reports subclonal CNAs, the values of RDR and BAF of the corresponding sample-subclonal clusters are not clearly distinguished and are generally similar to the ones of sample-clonal clusters (Fig. 3E-F). While it is possible that Battenberg has higher sensitivity in detecting *subclonal CNAs* than HATCHet, both methods infer similar fractions of the genome affected by CNAs on these 20 samples (Fig. S30A), suggesting that the subclonal CNAs inferred by Battenberg are clonal instead. To further quantify the differences in the inference of subclonal CNAs, we designed a metric called *clonality distance* to assess the presence of subclonal CNAs directly from the observed RDR and BAF. This metric supports HATCHet’s results on the absence/presence of subclonal CNAs (Fig. S52). Moreover, we note that Battenberg uses ≈6X more parameters than HATCHet on this dataset (Fig. S33), as it models the clonal composition of each segment independently from the others (Fig. S1). This substantially larger number of parameters raises the possibility of overfitting, and indeed Battenberg shows no improvement over HATCHet in explaining the observed RDR (Fig. S32). By modeling the dependency across segments and leveraging the global signals along the genome and across samples, HATCHet is able to reliably distinguish well-supported subclonal CNAs from noise in the data.

**Fig. 3:**
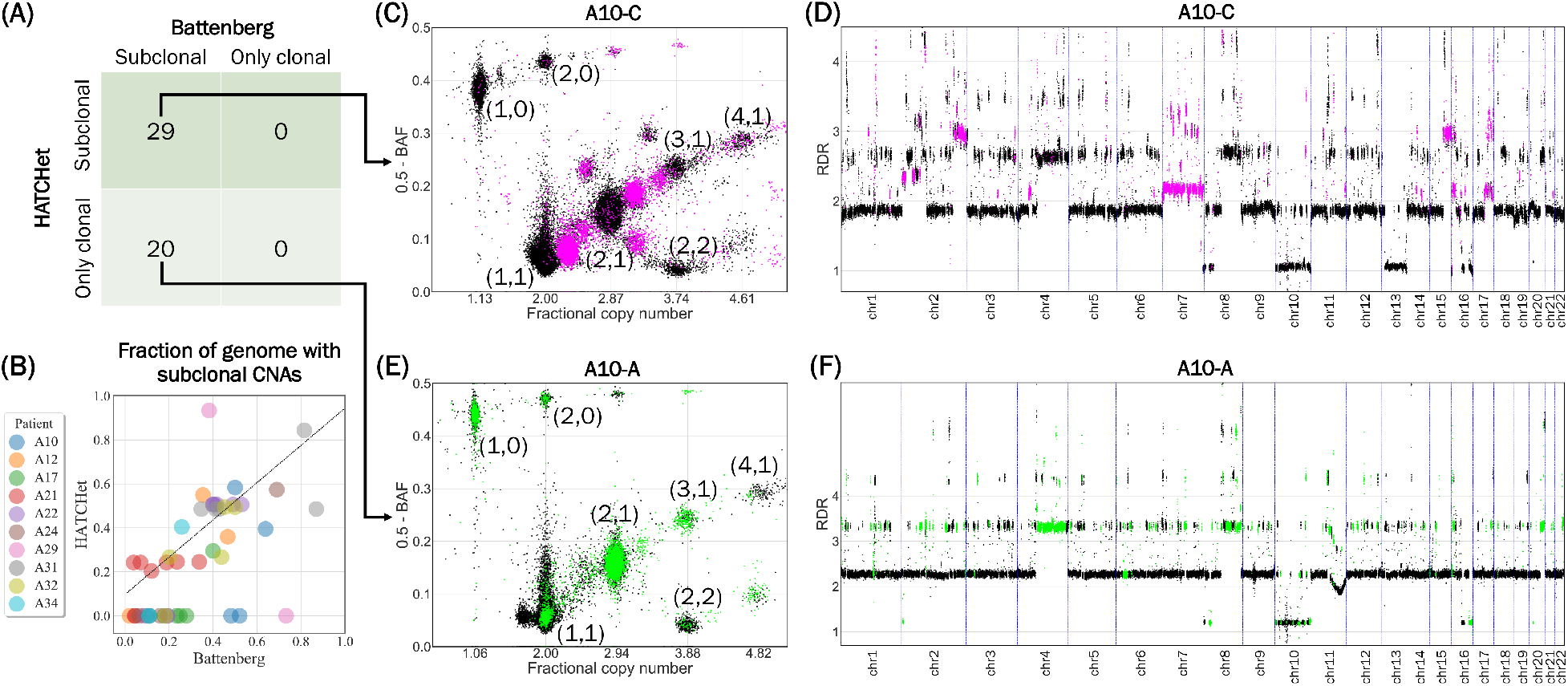
HATCHet identifies moderate amount of subclonal CNAs in prostate cancer patients. (A) HATCHet identifies subclonal CNAs in 29 samples, while Battenberg identifies subclonal CNAs in all 49 samples. (B) In the 29 samples where both methods identify subclonal CNAs, HATCHet and Battenberg infers similar fractions of the genome with subclonal CNAs (dotted line), while in the other 20 samples only Battenberg retrieves such a significant fraction. (C) In sample A10-C of patient A10, both HATCHet and Battenberg identify reliable subclonal CNAs that correspond to sample-subclonal clusters (magenta) with clearly intermediate positions in the scaled RDR-BAF plot between those of sample-clonal clusters (black clusters with corresponding copy-number states). (D) The sample-subclonal clusters in (C) correspond to large genomic regions (magenta) with values of RDR clearly distinct from the RDR values of regions from sample-clonal clusters (black). (E) In sample A10-A of patient A10, Battenberg identifies an extensive number of subclonal CNAs corresponding to sample-subclonal clusters (green). The green sample-subclonal clusters are not clearly distinguished from the clonal CNAs inferred by HATCHet (black clusters with corresponding copy-number states). (F) The sample-subclonal clusters in (E) correspond to large genomic regions (green) with values of RDR approximately equal to the RDR values of nearby regions from sample-clonal clusters (black).

In the pancreatic cancer dataset, HATCHet identifies subclonal CNAs in 20 of 35 samples (Fig. 4A). Published analysis used Control-FREEC for CNA inference, which does not consider the presence of subclonal CNAs, assuming instead that all CNAs are *clonal* and contained in all tumor cells in a sample. The sample-subclonal clusters inferred by HATCHet are well supported by the data with values of RDR and BAF that are clearly distinct from those of sample-clonal clusters (Fig. 4D) and correspond to whole-chromosomal arms (Fig. 4E). The clonality distance further supports the conclusion of subclonal CNAs in these samples (Fig. S53). Interestingly, many of the sample-subclonal clusters are also *tumor-subclonal*, meaning that these clusters do *not* have the same copy-number state in all tumor clones (Fig. 1F and Fig. S9). However, the distinct copy-number states are shared between samples (Fig. 4B-C-D), indicating the presence of shared clones with distinct CNAs (see Section 2.5). HATCHet’s global clustering enables the identification of this clonal composition even in low-purity samples (e.g. Pam01_LiM1 in Fig. 4B or Pam02_LiM5 in Fig. 6B) by leveraging the signals from high-purity samples (e.g. Pam01_LiM2 in Fig. 4C or Pam02_PT18 in Fig. 6B). Interestingly, the identification of the same clusters in low-purity samples would have not been possible with standard local clustering – which considers each sample independently – because distinct clusters are complicated to identify in low-purity samples. Overall, we find that in comparison to the published analysis (which did not consider the presence of subclonal CNAs), HATCHet reports a greater fraction of the genome with CNAs (Fig. S31) and better fits the observed RDR and BAF (Fig. S34) using fewer than 1/3 of the parameters used by Control-FREEC (Fig. S35).

**Fig. 4:**
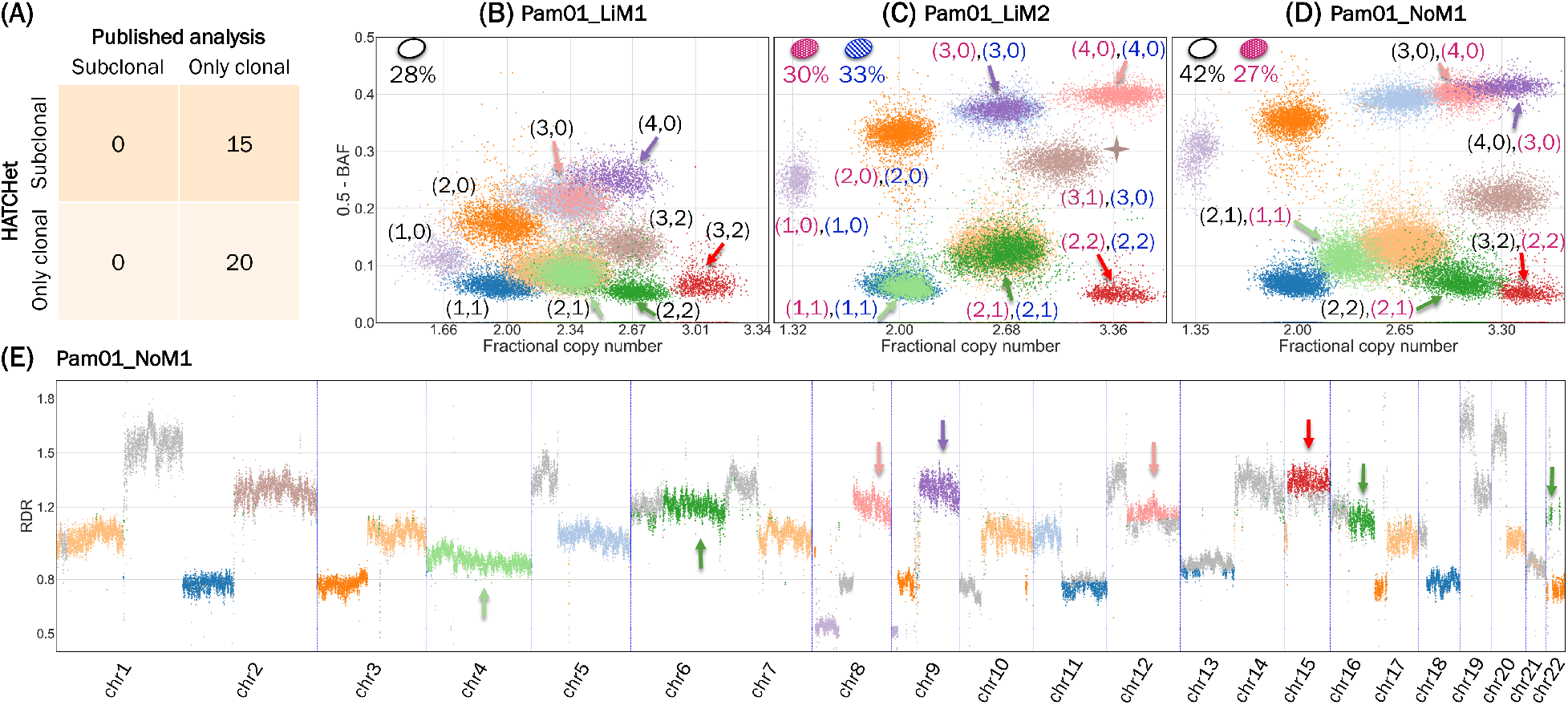
HATCHet identifies well-supported subclonal CNAs in metastatic pancreas cancer patients. (A) HATCHet identifies subclonal CNAs in 15 of 35 samples, while published analysis did not consider the presence of subclonal CNAs. (B) In the liver metastasis sample Pam01_LiM1, HATCHet identifies a single tumor clone and infers low tumor purity (28%). (C) In a second liver metastasis sample Pam01_LiM2 from the same patient, HATCHet identifies two distinct tumor clones (ellipses in upper right of plot with corresponding proportions) and infers higher tumor purity (63%). The largest sample-subclonal cluster (brown, starred) correspond to subclonal CNAs (i.e. distinct copy-number states in the two clones) and occupies an intermediate position between the other sample-clonal clusters in the scaled RDR-BAF plot. 5 tumor-subclonal clusters (arrows) have different copy-number states in the clones in (B) and (C) and thus vary their relative positions in the scaled RDR-BAF plots. (D) In a lymph node metastasis sample Pam01_NoM1 obtained from the same patient, HATCHet infers a mixture of the clone in (B) and one of the clones in (C). The 5 tumor-subclonal clusters (arrows) are subclonal in sample Pam01_NoM1 and the copy-number states associated with each of these clusters in this sample are mixtures of the different states associated with the corresponding clusters in Pam01_LiM1 and Pam01_LiM2. These sample-subclonal clusters are well supported by their large size, the high inferred purity of the sample (69%), and their intermediate positions in (D). The shared clones between the different metastases (C) and (D) suggest a crucial role of lymph nodes in the evolution of this tumor. (E) The 5 sample-subclonal clusters (arrows) correspond to large genomic regions and have clearly distinct values of RDR in Pam01_NoM1 with respect to the other sample-clonal clusters. Genomic regions that are part of small clusters or have out-of-scale values are reported in gray.

**Fig. 5:**
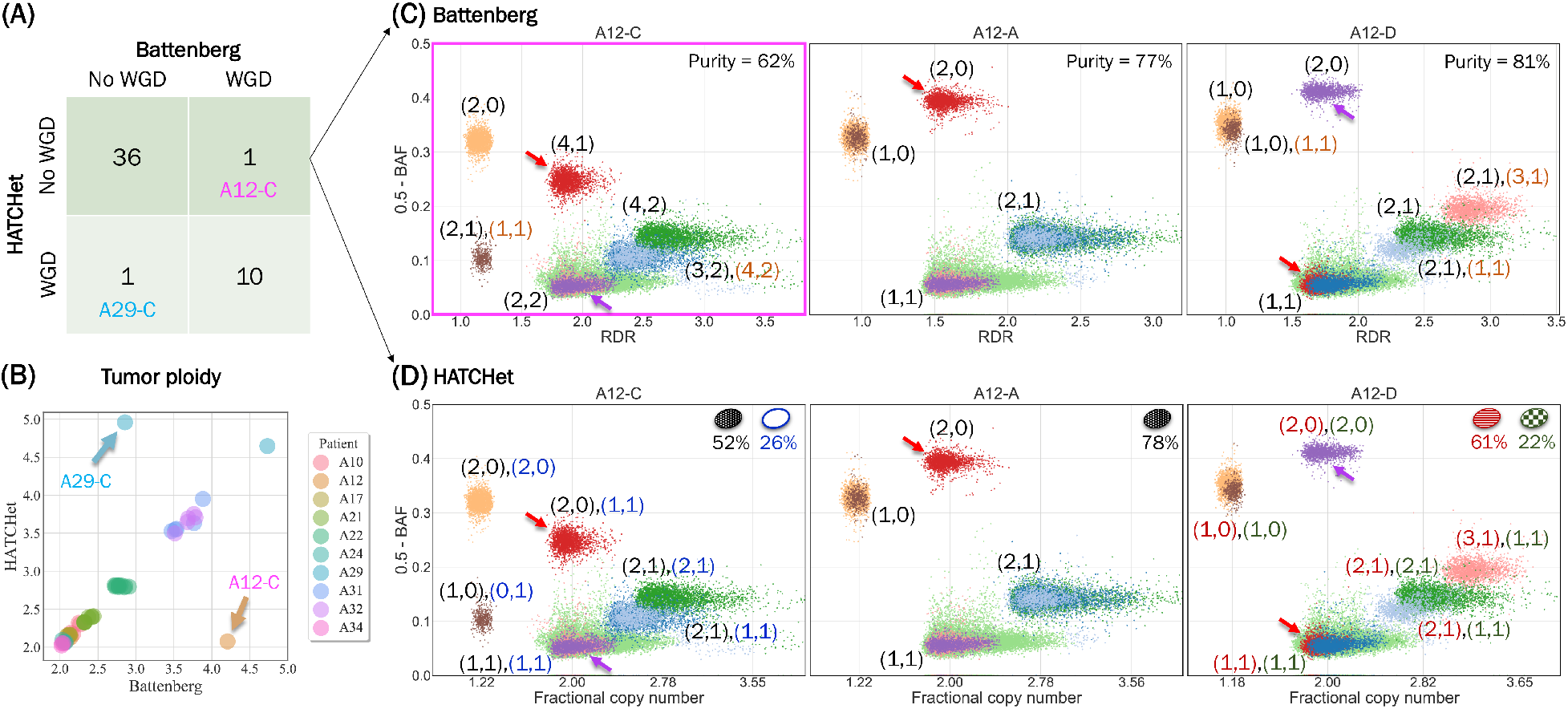
HATCHet predicts WGDs consistently across all samples from the same prostate cancer patient. (A) HATCHet predicts WGDs in 47 of 49 samples concordantly with the published analyses based on manual review of Battenberg’s tumor ploidy. While HATCHet predicts WGD consistently across all samples from the same patient, Battenberg predicts the presence of a WGD only in the sample A12-C of patient A12 and the absence of a WGD only in the sample A29-C of patient A29. (B) The tumor ploidies predicted by HATCHet and Battenberg are nearly identical in all samples except A12-C and A29-C. (C) The copy-number states inferred by Battenberg for three samples from patient A12, where a WGD was predicted only in sample A12-C. The larger number of clusters in A12-C could be explained by either the presence of subclonal CNAs or a WGD; however Battenberg infers *both*, even though it does not predict a WGD in the other two samples, A12-A and A12-D. The Battenberg’s solution is also unlikely because of the copy-number states inferred for the purple cluster. The WGD in A12-C cannot occur after the complete loss of one allele for the purple cluster in sample A12-D, as the lost allele cannot be re-acquired. Moreover, the WGD in A12-C is unlikely to have occurred first, as many of the clusters in A12-D would then have to revert to their pre-WGD state. Thus, the only plausible explanation is that the WGD and transition of the purple cluster from the (1,1) to the (2, 0) state occurred on different phylogenetic branches; however, even this explanation is unlikely, as other clusters in A12-D would also have to transition in a coordinated way on these parallel branches. Finally, the red and light-green clusters almost have the same RDR in A12-C but Battenberg infers different total copy numbers for these (4 vs. 5). (D) HATCHet does not predict a WGD in any sample from patient A12, instead inferring the mixture of two sublcones in samples A12-C and A12-D. Importantly, the red cluster is the *only* cluster in sample A12-C whose clonal/subclonal status differs from the Battenberg solution in (C). The position of the red cluster in the scaled RDR-BAF plot in A12-C is clearly intermediate between the positions of this cluster in other two samples (all with similar values of tumor purity), supporting HATCHet’s interpretation of the red cluster in sample A12-C as a mixture of the copy-number states (2, 0) and (1,1) of the red cluster in samples A12-A and A12-D, respectively.

**Fig. 6:**
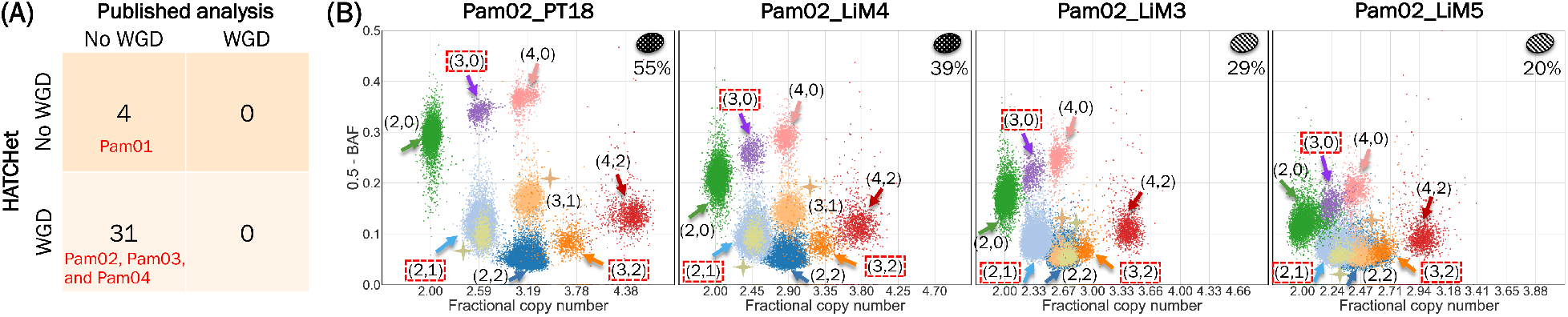
HATCHet identifies WGDs in three of four pancreas cancer patients. (A) HATCHet predicts a WGD in all 31 samples of 3 patients (Pam01, Pam02, and Pam03). In contrast, published analysis excludes WGDs. (B) In four samples of patient Pam02, HATCHet predicts a WGD and infers two tumor clones (ellipses in upper right of plot with corresponding proportions) with 7 large tumor-clonal clusters (arrows with corresponding copy-number states). These clusters preserve their relative positions across samples and their fractional copy numbers correspond to sample-clonal clusters in each sample (vertical grid lines) supporting the inference of a tumor-clonal CNA (i.e. unique copy-number state) for each of these clusters. Two additional clusters (peach and olive, starred) are tumor-subclonal as they change their relative position across samples (Pam02_PT18 and Pam02_LiM4 vs. Pam02_LiM3 and Pam02_LiM5), supporting the inference of two distinct tumor clones in this patient.

Additional details for all the results and analyses are reported in Supplementary Note C.2.

### 2.4 HATCHet reliably identifies whole-genome duplications

We next examine the prediction of whole-genome duplications (WGDs) on the prostate and pancreas cancer datasets. Battenberg does not explicitly state whether a WGD is present in a sample, and thus we use the criterion that tumor ploidy > 3 corresponds to WGD, following the values of tumor ploidy in previous pan-cancer analysis^5–8,12^. Using this criterion, there is strong agreement between WGD predictions from Battenberg and HATCHet on the prostate dataset, with discordance on only 2 of 48 samples (Fig. 5A,B). We note that Battenberg’s solutions were manually chosen from many alternatives, and the strong agreement between these predictions is thus a positive indicator for HATCHet’s automated model selection. The 2 discordant samples (A12-C and A29-C) are single samples from two different patients (A12 and A29, respectively); thus Battenberg predicts the presence of a WGD in only 1 of the 3 samples from patient A12 and the absence of a WGD in only 1 of the 2 samples from patient A29. This prediction of WGD in only a subset of tumor samples from a single patient is unlikely and not well-supported by the data from these patients. In general, a large number of copy-number states (i.e. large number of clusters) is a signal of either a WGD or subclonal CNAs (Fig. S3). However, Battenberg predicts *both* a WGD *and* the presence of subclonal CNAs in these 2 samples (Fig. 5C and Fig. S12A). In contrast, HATCHet infers WGD consistently across all samples from the same patient, and explicitly considers WGD in the model selection step. The result is simpler solutions that are consistent and well-supported across the multiple samples from the same patient (Fig. 5D and Fig. S12B). Moreover, HATCHet’s solutions are well supported by the clonality distance metric (Fig. S54A).

On the pancreas dataset, the published analysis excludes WGDs and assumes that tumor ploidy is always equal to 2. Instead, HATCHet predicts a WGD in all 31 samples of three of the four patients (Fig. 6A), and infers high tumor ploidy (> 3) for all samples from all four patients (Fig. S13A), even in the samples of patient Pam01 where HATCHet does not infer a WGD. These results are consistent with recent reports of the high frequency of WGD (~45%) and massive rearrangements in pancreatic cancer^26,43^. All 31 samples from the 3 patients with a WGD display a significant number of clusters of genomic regions with clearly distinct values of RDR and BAF. When jointly considering all samples from the same patient, these clusters are clearly better explained by the occurrence of a WGD (Fig. 6B) than by the presence of many subclonal CNAs, as the latter would result in the unlikely presence of two tumor clones with the same proportions in all samples (Fig. S13B). By directly evaluating the trade-off between subclonal CNAs and WGDs in the model selection, HATCHet makes more reasonable predictions of the occurrence of a WGD, which is also well supported by the clonality distance (Fig. S54B).

### 2.5 HATCHet enables the identification of tumor clones shared across samples

The published analysis of the prostate and pancreas cancer datasets reported that most SNVs are shared across samples from the same patient. This observation led to the conclusion that there is limited heterogeneity between the samples, with several samples sharing the same set of tumor clones^11,30^. However, the published CNAs generally do not support the reported limited heterogeneity. To quantify this discordance, we identified *sample-specific copy-number states*, i.e. copy-number states (*a*, *b*) that are unique to a single sample (Fig. S15A). We observe that the published CNAs for the prostate and pancreatic datasets contain sample-specific copy-number states in *every* sample in a significant fraction of the genome (Fig. S15B–C and Fig. S36–S37); these correspond to many, large sample-specific CNAs distributed across *all* chromosomes (Fig. S38 and Fig. S39). This is not surprising since the CNA methods used in these studies (Battenberg and Control-FREEC) identify CNAs separately in each sample. In contrast, HATCHet identifies sample-specific copy-number states in only a few samples, a consequence of HATCHet’s joint analysis of tumor clones across samples (Fig. S10 and Fig. S11). Overall, the CNAs inferred by HATCHet support the previously-reported limited heterogeneity across samples (especially metastatic samples) better than published CNAs.

A key finding in the prostate publication^11^ was the presence of subclonal clusters of SNVs that were shared across different primary and metastasis samples of 5 patients (A22, A24, A31, A32, and A34). This observation led the authors to conclude that some metastases in these patients were a result of *polyclonal migrations*. Curiously, the published CNAs support an even more complicated story, as Battenberg infers a significant amount of shared subclonal CNAs in *all* samples of the 10 patients, except those with different predictions of WGDs (Fig. S14). In contrast, the CNAs inferred by HATCHet confirm the presence of multiple CNA-derived tumor clones shared between multiple samples from only 3 (A22, A31, and A32) of the 5 patients (Fig. 7 and Fig. S16). Interestingly, these same 3 patients were also the only ones reported with polyclonal migrations in another analysis of this dataset using the MACHINA algorithm for computing parsimonious migration histories^13^.

**Fig. 7:**
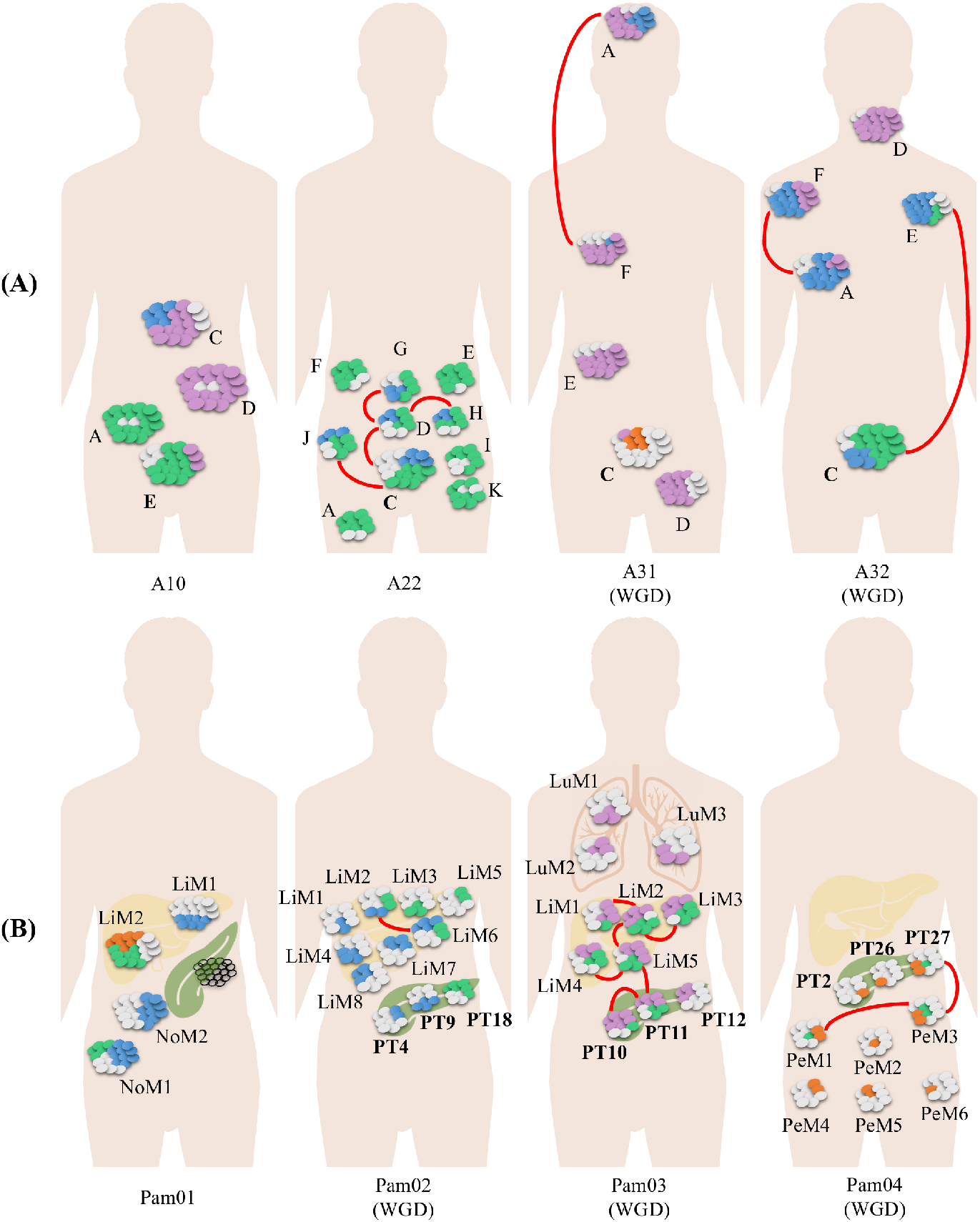
HATCHet identifies multiple tumor clones shared across samples from the same patient, suggesting polyclonal origin of metastasis in some prostate and pancreas cancer patients. HATCHet infers a normal clone (gray ellipses) and one or more tumor clones (ellipses with an identifying color for each clone) shared across the samples of every patient (proportions of ellipses approximate the inferred clone proportions). Bold sample(s) are from primary tumor; other samples are metastases. Red arcs connect samples with two or more shared tumor clones, evidence of potential polyclonal migrations between anatomical sites. Patients for which HATCHet predicts a WGD are labeled correspondingly. (A) The 3 prostate cancer patients (A22, A31, and A32) with multiple tumor clones shared between some samples (red arcs) are the same three patients that were inferred to have polyclonal seeding via the MACHINA algorithm^13^, and a subset of the 5 patients reported to have polyclonal seeding in the original published analysis^11^. (B) In pancreas cancer patient Pam01, lymph node metastasis sample NoM1 shares one tumor clone (blue) with a liver metastasis sample LiM1 and a different tumor clone (green) with a distinct liver metastasis sample LiM2, suggesting a role for lymph nodes in metastasis in this patient. The other 3 pancreas cancer patients (Pam02, Pam03, and Pam04) have multiple tumor clones shared between some samples (red arcs), evidence of potential polyclonal migrations between anatomical sites. Sharing of tumor subclones between anatomical sites was not considered in the original published analysis^30^.

The pancreas cancer publication^30^ did not describe shared subclones between different samples from the same patient. In contrast, the CNA-derived clones from HATCHet identify several such cases. For example, in patient Pam01 HATCHet identifies two clones in a lymph node metastasis sample Pam01_NoM1 (Fig. 4D), one of which is found in a liver metastasis (sample Pam01_LiM1 in Fig. 4B) and the other found in a different liver metastasis (sample Pam01_LiM2 in Fig. 4C). This result suggests a crucial role for the lymph node in the metastatic spread of this tumor, a finding consistent with standard models of metastasis^44^ but in contradiction to recent studies from other cancer types that suggest lymph nodes do not actively participate in the metastatic process^45,46^. HATCHet also identifies multiple tumor clones shared across the primary and metastasis samples of 3 patients (Pam02, Pam03, and Pam04) (Fig. 7 and Fig. S17). This suggests the possibility of polyclonal migrations in these samples, consistent with the reports of polyclonal migrations in mouse models of pancreas tumors^47^.

### 2.6 HATCHet better explains somatic point mutations

As an independent measure of the quality of the CNAs and proportions inferred by different methods, we evaluate the read counts of somatic point mutations, including single-nucleotide variants (SNVs) and small indels. We compare the observed variant allele frequency (VAF) of each mutation with the VAF that is predicted by the inferred copy-number states and proportions at the corresponding genomic locus. Specifically, we classify a mutation as *explained* when the predicted VAF is within a 95% confidence interval (CI) (according to a binomial model with beta prior^34,35^) of the observed VAF (Fig. 8A). On the prostate cancer dataset, we analyze an average of 10,600 mutations per patient (Fig. S40) and find that HATCHet has both significantly fewer non-explained mutations than Battenberg on the samples of all but 1 patient – where the difference is small – (Fig. 8B) and lower errors across all patients (Fig. S44). HATCHet explains most of the mutations with high values of VAF (Fig. S42), while the non-explained mutations mostly have smaller VAFs (Fig. S46). The latter suggests the presence of additional clones distinguished by SNVs that accumulated after CNAs, as previously reported in the prostate publication^11^.

**Fig. 8:**
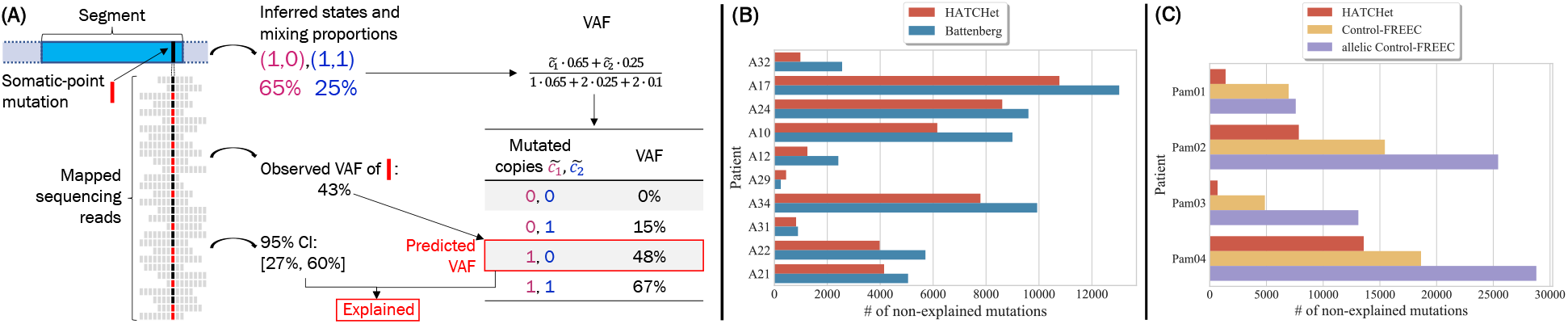
HATCHet infers copy-number states and proportions explaining somatic-point mutations better than published analysis. (A) A copy of a cyan genomic segment harbors a somatic-point mutation (red bar). For this segment, two copy-number states and relative proportions (corresponding colors) are inferred. From 30 sequencing reads covering that position (shifted sequences of bars), the observed variant-allele frequency (VAF) is computed as the fraction of reads harboring the mutation (red versus black bars) and the 95% confidence interval (Cl) on the VAF is obtained from a binomial model. If the number of mutated copies, 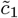 and 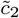, for each of the two copy-number states is known, then the VAF of the mutation is computed as the fraction of the mutated copies weighted by the proportions of the corresponding copy-number states. Assuming that an allele-specific position is mutated at most once during tumor progression (i.e. no-homoplasy), the *predicted VAF* is selected as the VAF that is closest to the observed VAF among the different possible values for the pair 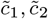. We say that the mutation is *explained* if the predicted VAF is within a 95% Cl of the observed VAF. (B) On the prostate dataset, HATCHet (red) explains more mutations than Battenberg (blue) in all patients but 1 (A29, where the difference is small), with the difference across all patients in excess of ≈ 10,500 mutations. (C) On the pancreas dataset, Control-FREEC does not provide allele-specific copy numbers. Thus, we assign the number of mutated copies by either considering both the alleles (yellow) or by first inferring allele-specific copy numbers according to the observed BAF of the segment (violet). In both cases and despite the bias of the former, HATCHet explains more mutations in all patients than Control-FREEC, with the difference across all patients in excess of ≈27,000 mutations and ≈ 56,100 mutations, respectively, in the two cases.

On the pancreas cancer dataset, we identify an average of 9,000 mutations per patient (Fig. S41) and also find that HATCHet has both significantly fewer non-explained mutations and lower errors than Control-FREEC on all four patients (Fig. 8C and Fig. S45). Moreover, we observe that nearly all mutations in the pancreas patients have low VAFs (Fig. S43). These low values can be explained either by low tumor purity of the samples or by the presence of WGDs and/or massive rearrangements. The latter events increase the copy numbers of most genomic regions and result, in general, in a lower proportion of copies harboring the mutations – particularly when the SNVs/indels occur after these events. Since we find that high-purity samples (e.g. Pam01_LiM2, Pam01_NoM1, and Pam02_PT18) also exhibit low VAFs, WGDs and high aneuploidy are more likely explanation, consistent with the copy numbers, WGDs, and proportions predicted by HATCHet (Fig. S47).

Finally, we assess whether the read counts of SNVs and small indels support the presence of multiple tumor clones in some samples, as inferred by HATCHet. To do this, we compute the cancer-cell fraction (CCF) – the fraction of tumor cells harboring a mutation – for each mutation using the copy-number states and proportions inferred by HATCHet and the number of mutated copies of the mutation inferred from the predicted VAF above. Explained mutations with a CCF less than 1 support the presence of more than one tumor clone in a sample (Fig. S48 and Fig. S49). We find that the tumor subclones inferred by HATCHet in both the prostate and pancreas datasets are supported by at least ≈2,000 SNVs on average (Fig. S50 and Fig. S51).

## 3 Discussion

The increasing availability of DNA sequencing data from multiple tumor samples from the same patient – including multiple regions of a primary tumor, matched primary and metastases, or longitudinal samples – provides the opportunity to improve the copy-number deconvolution of bulk samples into normal and tumor clones. Joint analysis of multiple tumor samples has proved to be of substantial benefit in the analysis of SNVs^11,29,32–36^. However, the advantages of joint analysis have not been exploited in the analysis of CNAs, with all analyses of the prominent multi-sample sequencing datasets (e.g.^11,12,29,30^) relying on CNA methods that analyze individual samples, and in some cases assuming that copy numbers are the same in all tumor cells in a sample.

The HATCHet algorithm introduced in this paper infers allele and clone-specific CNAs and the proportions of distinct tumor clones *jointly* across multiple tumor samples from the same patient. Moreover, HATCHet provides a fresh perspective on the copy-number deconvolution problem cutting through the barriers that limit the effectiveness of existing methods. First, the standard paradigm in CNA inference is to perform *segmentation* of read counts to leverage *local* correlations along the genome; however with multiple samples it is more effective to perform *clustering* of read counts *globally* along the genome and across samples. Second, the commonly used coordinates of *tumor purity* and *tumor ploidy* are ill-suited to summarize the subtleties of mixed copy numbers in a sample. We show that the identification of the diploid cluster (in the case of no WGD) or the copy numbers of two clusters (in the case of WGD) are sufficient to estimate fractional copy numbers. These fractional copy numbers enable the evaluation of different deconvolution solutions and the derivation of model selection criteria in the natural coordinates of copy number states and clone proportions.

Another issue slowing progress in copy-number deconvolution is the generation of simulated data that does not include multiple tumor clones in a mixture, does not account for the different genome lengths of tumor clones, and/or does not model WGD. To address these limitations and inaccuracies, we developed MASCoTE, a new simulator for multi-sample tumor sequencing data. On MASCoTE simulated multi-sample data, we demonstrate that HATCHet outperforms 4 current state-of-the-art methods (Battenberg, TITAN, THetA, and cloneHD), even when analyzing samples individually. Moreover, HATCHet significantly improves the identification of WGDs which has been reported to be common in many cancer types^5–7,12^. This improvement may enable the automatic prediction of WGDs on large cohorts of tumor samples, while current studies require manual inspection^5–7,12^ or rely on a consensus of biased methods^7^.

We demonstrate that HATCHet’s advantages result in simpler and more plausible inference of CNAs and WGDs on two whole-genome multi-sample tumor sequencing datasets. HATCHet’s inferred copy-number states contained a moderate number of subclonal CNAs and consistent WGDs. The CNAs and proportions inferred by HATCHet better explain the observed sequencing read counts of somatic SNVs in both datasets. Moreover, HATCHet identified shared CNA-derived clones between the samples from the same patient. These clones were generally more consistent with the SNV-derived clones in published analyses, and supported the previous reports of polyclonal origin of prostate metastases (in a subset of patients) and limited heterogeneity of metastasis in pancreatic cancer. Interestingly, the CNA-derived clones inferred by HATCHet suggest polyclonal origin of some pancreatic metastases.

While HATCHet is a substantial improvement over existing methods for CNA inference from bulk-samples, there are several areas for future improvements. First, while we have shown that HATCHet accurately recovers the major tumor clones distinguished by larger CNAs, HATCHet may miss small or minor CNAs, especially CNAs that are only present in a unique sample or in low proportions. One interesting direction is to perform a second stage of inference with a local segmentation algorithm informed by the clonal composition inferred by HATCHet. Second, HATCHet’s inference of WGD could be generalized further. Currently, we assume that at most one WGD occurs and that a WGD affects all tumor clones. Previous pan-cancer studies support these assumptions for most tumors^5–8,12^, but the reliable detection of subclonal WGD merits further investigation. Third, HATCHet might be improved by including other signals in DNA sequencing reads, including phasing of germline SNPs into haplotypes as in Battenberg^9^ or read counts of somatic point mutations. Fourth, the model-selection criterion of HATCHet could be extended to include additional parameters, such as the maximum copy number and the minimum clone proportion, and could be enriched with additional information such as the migration pattern between anatomical sites^13^. Fifth, a more refined model of copy-number evolution^23,24,48–50^ could be integrated in our model to simultaneously guide the factorization and obtain more information about the evolution and the temporal order of CNAs. Finally, some of the algorithmic advances in HATCHet can be leveraged in the design of better methods for inferring CNAs and WGDs in single-cell sequencing data.

The increasing availability of DNA sequencing data from multiple bulk tumor samples from the same patient provides the substrate for deeper analyses of tumor evolution across time and space, and in response to treatment. Algorithms that maximally leverage this data to quantify the genomic aberrations and their differences across samples will be essential in translating this data into actionable insights for cancer patients.

## 4 Method

We introduce HATCHet (<u>H</u>olistic <u>A</u>llele-specific <u>T</u>umor <u>Co</u>py-number <u>He</u>terogeneity), an algorithm to infer allele and clone-specific copy numbers and clone proportions for several tumor clones *jointly* across multiple bulk-tumor samples. We first formulate the copy-number deconvolution problem as a simultaneous matrix factorization problem. Next, we describe the 3 key components of HATCHet that extend the matrix factorization formulation into a practical algorithm for DNA sequencing data (Fig. 1): (1) global clustering of genomic regions along the genome and jointly across samples; (2) explicit estimation of fractional copy numbers; (3) an approach for addressing errors, uncertainty, and model-selection issues. Finally, we describe MASCoTE (<u>M</u>ultiple <u>A</u>llele-specific <u>S</u>imulation of <u>Co</u>py-number <u>T</u>umor <u>E</u>volution), a method to simulate DNA sequencing data from multiple bulk samples that correctly accounts for tumor clones with varying genome lengths.

### 4.1 Simultaneous matrix factorization model

We assume that each sample in our multi-sample tumor sequence set is a mixture of *n* clones. Each clone is distinguished by some number of copy-number aberrations (CNAs), where a CNA alters the number of copies of a contiguous genomic region from one of the two homologous chromosomes. We represent the accumulation of all CNAs in all clones by partitioning the *L* genomic positions of the reference genome into *m segments*, with each segment *s* consisting of *ℓ_s_* neighboring positions with the same copy number in every clone. Thus, a clone *i* is represented by a pair of integer vectors **a**_*i*_ and **b**_*i*_ whose entries indicate the number of copies of each of the two alleles for each segment. We define the *copy-number state*, or *state* for short, (*a*_*s*,*i*_, *b*_*s*,*i*_) of segment *s* in clone *i* as the pair of the two integer *allele-specific copy numbers a*_*s*,*i*_ and *b*_*s*,*i*_. We define *c*_*s*,*i*_ = *a*_*s*,*i*_ + *b*_*s*,*i*_ to be the *total copy number* of *s* in clone *i*. We define clone 1 to be the normal (non-cancerous) diploid clone, and thus (*a*_*s*, 1_, *b*_*s*, 1_) = (1,1) and *c*_*s*, 1_ = 2 for every segment *s* of the normal clone. We represent the allele-specific copy numbers of all clones as two *m* × *n* matrices *A* = [*a*_*s*, *i*_] and *B* = [*b*_*s*,*i*_]. Similarly, we represent the total copy numbers of all clones as the *m* × *n* matrix *C* = [*c*_*s*,*i*_] = *A* + *B*. Due to the effects of CNAs, the *genome length* 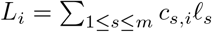 of every tumor clone *i* is generally different from the genome length *L*_1_ = 2*L* of the normal clone.

DNA sequencing data from a bulk tumor sample does not directly measure *A* and *B*, but rather measures a *mixture* of copy-number states. Specifically, each sample *p* is a mixture of clones, with *clone proportion u_i, p_* indicating the fraction of cells in sample *p* that belong to clone *i*. Note that 0 ≤ *u*_*i*, *p*_ ≤ 1 and the sum of clone proportions is equal to 1 in every sample *p*. We say that *i* is *present* in *p* if *u*_*i*, *p*_ > 0. Further, the *tumor purity* 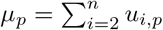 of sample *p* is the sum of the proportions of all tumor clones present in *p*. For a segment *s* from a sample *p*, we measure the *allele-specific fractional copy numbers* 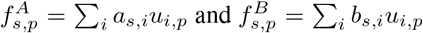 whose sum defines the *fractional copy number* 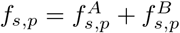.

The samples from a bulk-tumor are related by the somatic evolutionary process, and thus we model the fractional copy numbers *jointly* across the *k* samples from a tumor. Specifically, we represent the clone proportions as the *n* × *k* matrix *U* = [*u*_*i*, *p*_], and we represent the allele-specific fractional copy numbers using two *m* × *k* matrices 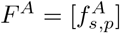 and 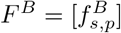. Then we have that *F^A^* = *AU* and *F^B^* = *BU*. The problem faced in bulk samples is to *simultaneously factorize F^A^* and *F^B^* into the corresponding allele-specific copy-numbers *A*, *B* and clone proportions *U* for some number *n* of clones. Formally, we have the following.

#### Problem 1

(Allele-specific Copy-number Factorization (ACF) problem). *Given the allele-specific fractional copy numbers F^A^ and F^B^ and the number n of clones, find allele-specific copy numbers A* = [*a*_*s*,*i*_], *B* = [*b*_*s*,*i*_] *and clone proportions U* = [*u*_*i*, *p*_] *such that F^A^* = *AU and F^B^* = *BU*.

The ACF problem differs in three key ways from current approaches to CNA inference:

1. ACF models allele-specific copy numbers, while many existing methods do not^20–24^.
2. ACF models dependencies *between samples* while all current approaches (with one exception^25^) analyze samples independently.
3. ACF models dependencies *between segments* as clones. Other widely-used methods either consider each segment independently^6,9,14–19^ (Fig. S1), do not preserve clonal structure across samples^21,22,26,27^, or assume all samples comprise the same set of few clones^25^.

While the ACF problem is a mathematically elegant description of the problem of inferring CNAs jointly from multiple mixed samples, there are several practical issues that must be addressed to derive a useful algorithm for DNA sequencing data:

1. The *m* segments that have undergone CNAs, which determine the entries of *F^A^*, *F^B^*, A, and B, are unknown.
2. *F^A^* and *F^B^* are not directly observed from DNA sequencing data.
3. Measurement errors in *F^A^* and *F^B^* may result in ACF not having any solution.
4. ACF is an underdetermined problem and multiple factorizations for given *F^A^* and *F^B^* may exist leading to degenerate solutions.
5. The number *n* of clones and the occurrence of WGD are unknown *a priori*.

In the following sections we describe how we address each of these issues.

### 4.2 Global clustering along the genome and across samples

The first practical issue is that genomic segments that have undergone CNAs in a sample must be inferred directly from sequencing-read counts. The standard approach to derive such segments is to assume that neighboring genomic loci with similar values of RDR and BAF are likely to have the same copy-number state in a sample. All current methods for CNA identification rely on such *local* information, and use segmentation approaches, such as Hidden Markov Models (HMMs) or change-point detection, to RDR and BAF measurements^9,14,15,17–19,51–53^.

With multiple sequenced samples from the same individual, one can instead take a different approach of identifying segments with the same copy-number state by clustering RDR and BAF *globally* along the genome and *simultaneously* across multiple samples (Fig. S2). Specifically, we use a non-parametric Bayesian clustering algorithm^54^ to cluster RDR and BAF values in short (≈ 50kb) genomic bins simultaneously across samples (further details are in Supplementary Note B.2). Each cluster thus corresponds to a set of segments with the same copy-number state in each tumor clone; these clusters can then be used to define the entries of *F^A^* or *F^B^*, playing the role of the segments described above. Although we do not require that clusters contain neighboring genomic loci, we find in practice that our clusters exhibit such locality (see results on cancer datasets). Thus, by clustering globally we preserve local information; but the converse does not necessarily hold. Moreover, the clustering of genomic regions in a sample with a low tumor purity is generally challenging because the variations in the values of RDR and BAF cannot be easily distinguished from noise in the data. However, the joint analysis on multiple samples leverages information from higher purity samples to assist in clustering of lower purity samples (see results on pancreatic cancer dataset).

### 4.3 Estimation of fractional copy numbers

In practice, one does not directly observe the allele-specific fractional copy numbers *F^A^* and *F^B^* from DNA sequencing data, but must infer these from the read counts of genomic segments. Widely used methods such as ABSOLUTE^6^, ASCAT^14^, Battenberg^9^, TITAN^17^, cloneHD^25^, and others^16,18–20,26,27^ do not attempt to directly infer fractional copy numbers, but rather attempt to fit other parameters, specifically the *tumor purity *μ*_p_* = 1 – *u*_1, *p*_ and *tumor ploidy* 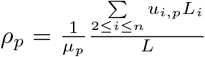 (or equivalent parameters as the haploid coverage, Supplementary Note B.1). However, tumor purity *μ_p_* and tumor ploidy *ρ_p_* are *composite* parameters that sum the contributions of the *unknown* copy numbers and proportions of multiple clones. This dependency is particularly complicated to model and easily becomes computationally challenging^21,22,25^. The consequence of this dependency is that tumor purity and ploidy are not good coordinates to evaluate tumor mixtures as many different clonal compositions may be equally plausible in these coordinates, particularly when more than one tumor clone in present. This ambiguity is especially prominent in the case of WGD as different values of *μ_p_* and *ρ_p_* can be equivalently inferred from the same read counts (Fig. S4) or the same values of RDR and BAF (Fig. S5). Not surprisingly, manual inspection of the results from current methods is often required to evaluate the presence of WGD^6,7,12^, while the few methods that attempt to automatize the prediction of WGD are based on biased criteria or unstated, restrictive assumptions^9,17,25^.

We introduce an approach to estimate *F^A^* and *F^B^* with rigorous and clearly-stated assumptions. First, in the case without a WGD, we assume there is a reasonable number of genomic positions in segments whose total copy number is 2 in all clones; this is generally true if a reasonable proportion of the genome is not affected by CNAs and, hence, is diploid. Second, in the case where a WGD occurs, we assume there are two groups of segments whose total copy numbers are the same in all clones and distinct; this is also reasonable if some segments are affected only by WGD and tumor clones accumulate common CNAs during tumor evolution. More specifically, we consider two signals obtained from the read counts of each segment *s* in every sample *p*: the read-depth ratio (RDR) *r*_*s*, *p*_ and the B-allele frequency (BAF) *β*_*s*, *p*_. Our approach scales *r*_*s*, *p*_ into fractional copy number *f*_*s*, *p*_ and separates *f*_*s*, *p*_ into the allele-specific fractional copy numbers 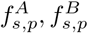 using 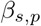. The following theorem states that the assumptions above are sufficient for scaling RDR to fractional copy numbers.

#### Theorem 1.

*The fractional copy number f_s, p_ of each segment s in each sample p can be derived uniquely from the RDR r_s, p_, *and either (1) a diploid clonal segment* s′, with total copy number c_s′,i_* = 2 *in every clone i or (2) two clonal segments s′,z′ with total copy numbers c_s′, i_* = *ω_s′_*, *c_z′, i_* = *ω_z′_ in every tumor clone i such that r_s′, p_*(*ω_z′_* − 2) ≠ *r_z′, p_*(*ω_s′_* − 2) *for all samples p*.

Notably, this theorem states that the scaling is independent of other copy numbers in *A*, *B*, and *C* as well as the clone proportions in *U*.

To apply this theorem, we design a heuristic for HATCHet to identify the required segments and their total copy numbers; our heuristic leverages the RDR and BAF jointly across multiple samples. First, in the case of no WGD, we aim to identify diploid segments with a copy-number state (1,1). These segments are straightforward to identify since *β*_*s*, *p*_ ≈ 0.5 across all samples *p* and we expect that a reasonable proportion of the genome in all samples will be unaffected by CNAs and thus have state (1,1). Since the total copy number of these segments is 2, these segments are sufficient to apply Theorem 1. In the case of a WGD, we assume that at most one WGD occurs and that any WGD affects all tumor clones, assumptions which are consistent with most tumors in previous pan-cancer analysis^5–8,12^. Our heuristic evaluates the same segments with *β*_*s*, *p*_ ≈ 0.5 as above, but now expect these to be tetraploid with a copy-number state (2, 2) as a WGD doubles all copy numbers. Since these segments have total copy number of 4, they are not sufficient to apply the first condition of Theorem 1. Thus, we use the second condition of the theorem and aim to find another group of segments with the same state in all tumor clones. More specifically, HATCHet finds segments whose RDR and BAF in *all* samples indicate copy-number states that result from single-copy amplifications or deletions occurring before or after a WGD^5^; for example, copy-number state (2,0) is associated to a deletion occurring before a WGD while copy-number state (2,1) is associated to a deletion occurring after a WGD. Moreover, we select only those group of segments whose RDR and BAF relative to other segments is preserved in *all* samples; such preservation indicates that the copy-number state is fixed in all tumor clones (Fig. 1F). Further descriptions of Theorem 1 and this heuristic are in Supplementary Note B.3.

### 4.4 Measurement errors and model selection

The final issues to adapt the ACF problem to DNA sequencing data are: (3) addressing errors and uncertainty in the fractional copy numbers *F_A_* and *F_B_* resulting from their estimation from RDR and BAF; (4) ACF is an underdetermined problem with degenerate solutions, an issue that is further complicated when there are errors in *F*; (5) the number *n* of clones and the occurrence of WGD are generally unknown *a priori*.

To address the first issue, we do not solve the simultaneous factorization *F^A^* = *AU* and *F^B^* = *BU* exactly, but rather minimize the distance between the estimated fractional copy numbers *F^A^* and *F^B^* and the factorizations *AU* and *BU*, respectively, weighted by the corresponding size of the clusters. In particular, we define the distance 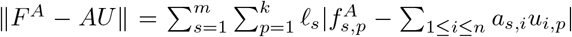, where *ℓ_s_* is the genomic length of the cluster *s*. We also define the corresponding distance for *F^B^*, *B*, and *b*.

To address the issue that the ACF problem is underdetermined with multiple degenerate solutions we include several reasonable constraints. First, since we do not expect copy numbers to be arbitrarily high – especially for large genomic regions – we constrain the simultaneous factorization by assuming that the total copy numbers are at most a value *c*_max_. Second, to avoid overfitting errors in fractional copy numbers by clones with low proportions, we require a *minimum clone proportion u*_min_ ∈ [0,1] for every tumor clone present in any sample. Third, we impose an evolutionary relationship between the tumor clones requiring that each allele of every segment *s* cannot be simultaneously amplified and deleted in distinct clones; i.e. either *a*_*s*, *i*_ ≥ θ *or a*_*s*, *i*_ ≤ θ for all clones *i*, where θ =1 when there is no WGD and θ = 2 when there is a WGD. The same constraint also holds for *b*_*s*, *i*_. We previously showed in^23,24^ that constraints based on the evolutionary process of CNAs may improve results for a related copy-number factorization problem. These evolutionary constraints are optional and less restrictive than the ones usually applied in current methods which, for example, assume: that tumor clones have at most two copy-number states per segment and the difference between allele-specific copy numbers is at most 1, i.e. |*a*_*s*,*i*_ − *b*_*s*,*i*_| ≤ 1 (as in^9,19^); or all clones have either a diploid copy-number state (1,1) or a unique aberrant state (*a*, *b*) = (1,1) in every cluster *s* (as in^17,18^); or every tumor clone *i* has either *c*_*s*, *i*_ ≥ 2 or *c*_*s*, *i*_ ≤ 2 for every cluster *s* (as in^21,22^); or there always exist segments with total copy number equal to 2 (as in^18,25^). We thus have the following problem.

#### Problem 2

(Distance-based Constrained Allele-specific Copy-number Factorization (D-CACF) problem). *Given the allele-specific fractional copy numbers F^A^ and F^B^; a number n of clones; a maximum total copy number c_max_, and a minimum clone proportion u_min_, find allele-specific copy numbers A* = [*a*_*s*,*i*_], *B* = [*b*_*s*,*i*_] *and clone proportions U* = [*u*_*i*, *p*_] *such that: the distance D* = ||*F^A^* + *AU*|| + ||*F^B^* + *BU*|| *is minimum; a*_*s*,*i*_ + *b*_*s*,*i*_ ≤ *c_max_ for every cluster s and clone i; either u*_*i*, *p*_ ≥ *u_min_ or u_i, p_* = 0 *for every clone i and sample p; for every cluster s, either a*_*s*,*i*_ ≥ *θ* or *a*_*s*,*i*_ ≤ *θ for all clones i; for every cluster s, either b*_*s*, *i*_ ≥ *θ* or *b*_*s*, *i*_ ≤ *θ for all clones i*.

We derive a coordinate-descent algorithm to solve this problem inspired by the algorithm we introduced in^23,24^ and we also derive an exact ILP formulation for small instances. Further details of this problem and methods are in Supplementary Note B.4.

Finally, we define a model-selection criterion to jointly select the number *n* of clones and predict the occurrence of a WGD. In general, variations in *F* can be fit by increasing the total number *n* of clones, increasing the number of clones present in a sample, or introducing additional copy-number states in a sample by inferring subclonal CNAs or WGD. There is a trade-off between these three options; for example, a collection of clusters exhibiting many different copy-number states may be explained by adding more clones (increasing *n*) and marking some clusters as subclonal or by the occurrence of a WGD with a larger number of clonal copy-number states (Fig. S3). Existing methods either: do not perform model selection and assume that the number *n* of clones is known^21,22,26,27^; ignore the trade-offs by considering segments independently^6,9,17–19^ (Fig. S1), perhaps increasing the sensitivity to detect small subclonal CNAs, but at the expense of overestimating *n* and the number of subclonal CNAs; ignore the specific trade-off between subclonal CNAs (related to a higher number of clones) and WGD by not including the presence of WGD in the model selection; perform model selection in the coordinates of tumor purity *μ_p_* and tumor ploidy *ρ_p_*, which does not adequately account for the ambiguity between different solutions as described above in Section 4.3.

Our model selection procedure consists of two steps. First, we observe that the distance *D* is a monotonically decreasing function of the number *n* of clones (following from^23,24^). Under the assumption that no WGD has occurred, we compute the distance *D* using the no-WGD-scaled *F_A_* and *F_B_* and find the value 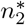 where the decrease in *D* first becomes small using standard “elbow criterion” from clustering. Similarly under the assumption that a WGD occurred, we compute the distance *D* using the WGD-scaled *F_A_* and *F_B_*, and find the value 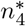 where the decrease in *D* first becomes small. If 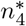 is smaller, we select this solution and infer a WGD. Otherwise, we select the solution with 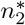 clones and no WGD. Our model-selection criterion differs from existing methods in two crucial points. First, our criterion explicitly examines the trade-off in number of clones (introducing subclonal CNAs) and the presence/absence of WGD while fitting the data to underlying parameters *A*, *B*, and *U*. In contrast, existing methods do not consider presence/absence of WGD during model selection and instead select a solution based on values of tumor purity and tumor ploidy, both of which are averages over the distinct tumor clones. Second, our criterion is based on an objective function that is monotonic in the parameter *n* and thus is better suited for model selection. This is in contrast to existing methods which attempt to fit the data using composite parameters of tumor purity and tumor ploidy, where model selection is complex due to non-monotonicity of distance/likelihood functions. Additional details of the model selection are in Supplementary Note B.5.

### 4.5 Simulation of bulk tumor sequencing data

The simulation of DNA sequencing data from bulk tumor samples that contain large-scale copy number aberrations is not straightforward, and subtle mistakes are common in previously published studies. Suppose *R* sequencing reads are obtained from a sample consisting of *n* clones with clone proportions *u*_1_,…, *u_n_*. Assuming that reads are uniformly sequenced along the genome and across all cells, what is the expected proportion *v_i_* of reads that originated from clone *i*? Most current studies that simulate sequencing reads from mixed samples (e.g.^15–17,25,37–42^) set *v_i_* = *u_i_*, although this selection is sometimes obscured in equations of slightly different approaches that use various poorly motivated and incorrect normalizations^1^ of *u_i_*. However, *u_i_* is the correct proportion *only* when the genome lengths of all clones are equal, i.e. *L_i_* = 2*L* for every clone *i*. Using an incorrect proportion *v_i_* leads to incorrect simulations of read counts, particularly in samples containing WGDs or multiple large-scale CNAs in different clones (Fig. S6 and Fig. S7). In fact, read counts depend on the genome lengths of *all* clones in the sample^55^ and the correct proportion 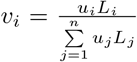 is equal to the fraction of genome content in a sample belonging to the cells of clone *i*. Moreover, the expected proportion *v_s, i_* of reads in segment *s* that originate from clone *i* is equal to 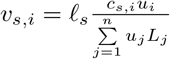, the fraction of the genome content from segment *s* belonging to the cells of clone *i* (Supplementary Note B.1).

To address these issues, we develop MASCoTE (<u>M</u>ultiple <u>A</u>llele-specific <u>S</u>imulation of <u>Co</u>py-number <u>T</u>umor <u>E</u>volution) to simulate sequencing data of multiple mixed samples obtained from the same patient (Fig. S8). MASCoTE simulates the genomes of a normal clone and *n* − 1 tumor clones which accumulate CNAs and WGDs during tumor evolution; these clones are related via a phylogenetic tree. As such, every sample comprises a subset of these clones and the corresponding sequencing reads are simulated according to the genome lengths and proportions of the clones. More specifically, MASCoTE is composed of four steps (Fig. S8): (1) MASCoTE simulates a diploid haplotype-specific germline genome (Fig. S8A); (2) MASCoTE simulates the genomes of *n* − 1 tumor clones that acquire different kind of CNAs and WGDs – according to the distributions in size and quantity reported in previous pan-cancer analysis^5^ – in random order through a random phylogenetic tree (Fig. S8B); (3) MASCoTE simulates the sequencing reads from the genome of each clone through standard methods^56^ (Fig. S8C); (4) MASCoTE simulates each sample *p* by considering an arbitrary subset of the clones (always containing the normal clone) with random clone proportions and by mixing the corresponding reads using the read proportion 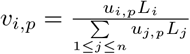 (Fig. S8D). Further details about this procedure are in Supplementary Note B.6.

## Acknowledgments

We thank Christine Iacobuzio-Donahue and Alvin Makohon-Moore for assistance in obtaining the copy-number data from their publication^30^. We thank Stefan Dentro, Peter Van Loo, and David Wedge for assistance in running Battenberg on our simulated data. We thank Gavin Ha for assistance in running TITAN on our simulated data. This work is supported by a US National Institutes of Health (NIH) grants R01HG007069 and U24CA211000 and US National Science Foundation (NSF) CAREER Award (CCF-1053753) to BJR.

## Code availability

HATCHet is available on GitHub at https://github.com/raphael-group/hatchet. MASCoTE is available on GitHub at https://github.com/raphael-group/mascote.

## Data availability

The prostate and pancreas cancer datasets analyzed in this study are available from the European Genome-phenome Archive (EGA) under accession numbers EGAS00001000262 and and EGAS00001002186, respectively. All the processed simulated data, the results of all methods on simulated data, and the results of HATCHet on the prostate and pancreas cancer datasets are available on GitHub at https://github.com/raphael-group/hatchet-paper.

## Footnotes

1 For example,^17,37^ artificially form a mixed sample of two clones by mixing reads from two other given samples in proportions 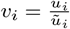 where *ũ_i_* is the clone proportion of the single tumor clone uniquely present in a given sample *i*. Another example is^40^ that simulates the reads for each segment *s* separately by setting 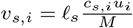 for every clone *i* where 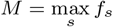 is the maximum fractional copy number.

